# Event-Related potentials in an associative word pair learning paradigm

**DOI:** 10.1101/741728

**Authors:** Maryam Farshad, Yuri G. Pavlov, Boris Kotchoubey

**Affiliations:** Institute of Medical Psychology and Behavioral Neurobiology, University of Tübingen, Germany; Department of Psychology, Ural Federal University, Ekaterinburg, Russian Federation

**Keywords:** Associative learning, ERP, Late Positive Complex (LPC), N400, Priming

## Abstract

The study investigated the effect of unintentional learning of semantically unrelated word pairs on event-related brain potentials. Two experiments were conducted, in whose acquisition phase participants listened to five pairs of semantically unrelated words, each pair being repeated twenty times. In the test phase of Experiment 1, these “old” pairs were presented mixed with “new” pairs containing other words. In the test phase of Experiment 2, a third condition was added in which the first word in a pair was one of the words presented during acquisition but the second word was new. In both experiments, the second word in new word pairs elicited an N400 and a late (550-1000 ms) frontal positivity. The amplitude of the N400 to new second words in Experiment 2 was significantly larger when the first word in the pair was an old (previously learnt) word, as compared with the condition in which both first and second words were new. The results indicate that, in addition to a repetition effect, unintentional learning of word pairs results in building new associations between previously unrelated words.

## 1. Introduction

Investigation in the electrophysiology of semantic processing began with a famous study of Kutas & Hillyard (1980) who showed that a negative deflection peaking around 400 ms post-stimulus strongly distinguished between the last words in sentences that were semantically congruent versus incongruent with the context of the sentence (e.g., “I drink my coffee with cream and *sugar*” versus “I drink my coffee with cream and *dog*”). This deflection, labeled N400, was shown to be a part of the normal brain response to words and other meaningful (or potentially meaningful) stimuli but it is typically attenuated when the word is strongly prepared by its context (Kutas & Federmeier, 2011).

In addition to the above-depicted sentence paradigm with congruent versus incongruent endings, similar results were obtained in experiments with word pairs (Holcomb, 1988). In both paradigms, the response to a word is regarded as a function of the background, or the context, on which the word appears (Weingarten et al., 2016). In the word pair paradigm, the context for the second word in a pair (referred to as “target”) is built by the first word (“prime”). In the sentence paradigm, the last word is primed by the context of the sentence. N400 amplitude is typically larger for targets that are incongruent with their primes than for congruent targets and inversely related to the degree of semantic expectancy. The similar paradigm can be extended to an even larger context of the whole conversation (e.g., van Berkum, Hagoort, & Brown, 1999): N400 amplitude is increased in response to targets incongruent with the idea of the conversation, even if they are completely congruent with the immediately preceding sentence. Generally, the modulation of the N400 amplitude by the preceding context is referred to as the semantic priming effect (Brualla, Romero, Serrano, & Valdizán, 1998).

In addition to the semantic relationship, an association between two words can emerge due to their spatial and/or temporal co-occurrence in the real world and/or in a language (McRae, Khalkhali & Hare, 2012). N400 effect is sensitive to how often two concepts co-occurred within an individual’s past experience. An example of such co-occurring concepts is the pair rock and roll (Lancia, 2007; Buchanan, Holmes, Teasley & Hutchison, 2013). One word can remind us of the other one because of their main association with the music style, although there is no apparent semantic relationship. Moreover, there is evidence of a graded effect of association strength: the N400 to the second word in moderately associated word pairs (e.g., protection – cover) is larger than in strongly associated pairs (e.g., atom – bomb) but smaller than in the pairs without association (e.g., parade – slice) (Ortu, Allan and Donaldson, 2013).

While numerous studies have investigated the effects of associations learnt during the previous life, only a few works were devoted to associative relationships built immediately in the course of the experiment. In the acquisition phase of the experiment of Balass, Nelson and Perfetti (2010), participants learned very rare English words. Specifically, the learned either to associate spelling of the word with its pronunciation (OP condition), or spelling with meaning (OM condition), or pronunciation with meaning (PM condition). In the test phase word pairs were presented, whose first word was either one of the just learnt rare words, or a new rare word, or a highly familiar word. The N400 to the second word was larger to new rare words and to words previously learnt in the OP condition that to highly familiar words and to rare words learnt in the OM or PM condition. Thus, the results indicated that the N400 is specifically affected by the semantics of newly learnt words rather than their orthography and pronunciation.

The data of another semantic learning experiment (Borovsky, Kutas and Elman, 2010) lead to the same conclusion. In the leaning phase of the experiment, participants were presented real words or pseudowords in sentences strongly or weakly restraining their meaning (examples for a pseudoword being: He tried to put the pieces of the broken plate back together with MARF, and She walked across the room to Mike’s desk to return his MARF, for strong and weak context, respectively). In the test phase participants had to decide whether a sentence had an appropriate (e.g., they used a MARF) or inappropriate (e.g., they greeted a MARF) meaning. The most interesting result was obtained in response to the verb (“used” or “greeted”) in the test sentence: for real words regardless of the context in the learning phase, and for pseudowords presented with strong context (but not for pseudowords presented with weak context), the N400 was significantly larger to inappropriate than appropriate verbs.

Sometimes, the N400 in response to verbal stimuli is followed by a late positive component (LPC) – a long-latency, positive-going shift, usually with a parietal maximum. Although semantic learning can occasionally happen with a single presentation (see Borovsky et al., 2010), mostly words are presented repeatedly to learn. The main effect of repetition of words in lists is the marked enhancement of the LPC (e.g., Rugg, 1985; Rugg, Pearl, Walker, Roberts, and Holdstock, 1994; Rugg, Doyle, & Holdstock, 1994) – either along with a reduction of the N400 or without it (Van Petten, Kutas, Kluender, Mitchiner and McIsaac, 1991). As compared with the semantic N400 effect described above, the similar N400 effect elicited by word repetition peaks later and lasts longer (e.g., Besson et al., 1992).

Whereas the N400 priming effect contrasting semantically related versus semantically unrelated word pairs has been carefully investigated in numerous experiments (e.g., Holcomb, 1993; Kiefer, 2005; Kiang et al., 2013), we could not find any study that contrasted two groups of equally unrelated word pairs: learnt versus unlearnt pairs. Since learning of non-associated word pairs involves their repeated presentation, a repetition effect can be expected. However, in the current case two repetition effects can be dissociated: (1) single word level repetition effect that may cause similar effects of N400 reduction with an increase of the LPC, and (2) the repetition of the association within word pairs, that has not been studied by previous research.

In two experiments we tested whether unintentional learning of semantically not associated word pairs would result in N400 attenuation similar to that in semantically associated pairs (Holcomb, 1993; Balass et al., 2010). We compared ERPs to unrelated word pairs that have been recently learnt (“old” pairs) and to other unrelated word pairs that were presented for the first time (“new pairs”). In the first experiment, we expected that the N400 to the second word in a pair to be smaller in learnt word pairs than in new word pairs because consistent repetition of semantically unrelated word pairs may make them acting like related word pairs. The effect is expected to be stronger than the repetition effect on the first words, thus indicating that associated pair repetition involves some processes beyond simple word repetition. Further, we aimed to clarify whether the N400 effect is to be explained by increased familiarity of repeatedly presented words or is based on building new associations between the members of a pair. For this sake, in the second experiment we introduced so-called old-new where the first word was already learnt in another association, and the second word was new (unlearnt). We hypothesized that, if the new association between words in pairs has an effect independent of that of repetition, then the N400 to the second word in the old-new pairs would have larger amplitude as compared to the new-new pairs.

## 2. Methods

### 2.1 Participants

Twenty-two subjects participated in this study. The group comprised thirteen females and nine males with a mean age of 25.23 (range: 19-33, standard deviation: 12.45) recruited from the University of Tuebingen student population. None of them reported visual, auditory or neurological deficits. According to the Edinburgh Handedness Scale (Oldfield, 1971), seventeen participants were right-handed, four left-handed and one participant was ambidextrous. They were either native German speakers or came to Germany before the age of 6 and started their primary school here. Informed consent was obtained from all subjects included in the study. They received an expense allowance of 8 €/h for their participation. One participant’s data were excluded from some of the analyses because of missing data in the acquisition phase of the second experiment due to EEG recording failure.

### 2.2 Stimuli and procedure

We carried out two learning experiments, each entailing two phases. We hypothesized that the result of learning in the first (acquisition) phase should be manifested in the second (test) phase.

Experiment 1 contained an acquisition phase, in which five word pairs were presented 20 times each, ten times in one order (e.g., warm / fast) and ten times in the reverse order (e.g., fast / warm). In the test phase of Experiment 1 the same five word pairs from the acquisition phase were presented (“old” pairs) 20 times each, resulting in 100 trials, and there were 100 trials with 50 completely new word pairs, each of which was presented twice (once in one order and once in the reverse order). None of these words had been presented during acquisition.

The acquisition phase of Experiment 2 was identical to the acquisition phase of Experiment 1. The test phase included 300 word pairs subdivided into three categories: 100 “old” pairs presented at the acquisition phase and 100 new pairs were analogous to those in Experiment 1. An additional category involved 100 word pairs, whose first word was the same as one of the “old” words, learnt in the acquisition phase. However, the second word in the pair was new. These pairs were designated as “old-new pairs”. None of the words (“old” or “new”) employed in Experiment 2 repeated any word from Experiment 1 (see Appendix for the full list of words). Absolute word count within the Deutsches Referenzkorpus (Kupietz, Belica, Keibel, & Witt, 2010) was (mean ± standard deviation) 274363±291642 and 480902±1191510, for old and new words, respectively (t(218) = 0.77, p = .441, d = −0.18); number of syllables was 1.3±0.470 and 1.25±0.459, for old and new words, respectively (t(218) = 0.42, p = .677, d = 0.10); word length during speaking (in milliseconds) was 664±103 and 667±104, for old and new words, respectively (t(218) = 0.11, p = .79, d = −0.026). Thus, there were no significant differences between old and new pairs regarding the basic characteristics of words. The two versions of the acquisition-test word pair sets (used in Experiment 1 and Experiment 2) were counter-balanced between participants.

The two above-described EEG experiments were preceded by a pilot paper-and-pencil experiment with fifteen students of medicine and psychology (nine females, all native German speakers). They were presented with 1320 different word pairs and were required to judge the association strength within each pair using the scale from 0 (no association at all) to 8 (very strong association). Moreover, they were instructed that they should regard word pairs such as man/woman and cat/mouse as “8”. After this, only word pairs whose association strength was never assessed above 3 were employed in the two EEG experiments.

In both experiments, the participants were sitting in a comfortable chair with the eyes closed. The words were presented auditorily by a female voice, without local accent. The loudness was normalized to be about 65 dB SPL. The word pairs were presented in a pseudorandom order, and one old pair was never presented twice in a row. Furthermore, in the test phases not more than four pairs of the same category (i.e., old, new, old-new) were presented in immediate succession. The interval between the offset of the first word and the onset of the second word within each pair was 100 ms. Due to the variable length of the words, the onset-to-onset interval varied from 500 to 1100 ms. The second word was followed by a motor response (see below), and the next word pair started 1 s after the motor response. In each experiment, the test phase started immediately after the end of the acquisition phase without a break. There was a break of about 30 minutes between the two experiments, during which participants were busy activities unrelated to the experiments.

Participants were asked to compare the number of syllables in the two words of each pair. If both words had the equal number of syllables, participants had to press one of the keyboard buttons (left or right Ctrl buttons on a standard keyboard), and if they had a different number of syllables, the other button. The side of the response was counterbalanced among the participants. The task was irrelevant concerning the aim of the experiments. It was just used to increase attention to the words. Because the mean length of experiment 1 was around 19 minutes and experiment 2 was about 27 minutes, we were concerned that participants’ attention would decline if no overt response is required.

### 2.3 EEG recording and ERP analysis

The EEG was recorded from 64 channels online referenced to Cz. The EEG was acquired with 1000 Hz sampling rate with a low-pass 280 Hz filter. The active electrodes (EasyCap) were placed using the 10-10 system. BrainVision Analyzer Version 2.1.2.327 was used for preprocessing of EEG. First, the data were downsampled off-line to 256 Hz and then re-referenced to a common average reference. After that, offline low pass 30 Hz and 0.1 Hz high-pass filters were applied. Bad channels were replaced using spherical interpolation. An Independent Component Analysis (ICA) was employed to separate and remove activity related to ocular artifacts. After artifact correction four channels (Fp1, Fp2, AF7, and AF8) were removed from further analyses. Finally, the data were again re-referenced to average mastoids to warrant the comparison with the published results, most of which have been reported with mastoid reference. ERPs were averaged in epochs started −200 ms before a stimulus and lasted for 1000 ms after it. The baseline was defined as the average amplitude between −200 ms and 0 ms. The epochs still containing artifacts were identified and rejected from further analysis by automatic artifact rejection algorithm implemented in BrainVision Analyzer.

The mean ERP amplitude was measured in the time windows of 250-550 ms (N400) and 550-1000 ms (LPC). We did not observe a clear peak in the grand average ERPs after N400 time window; hence we defined LPC from the end on N400 window to the end of the epoch. The statistical analysis was conducted using afex package (Singmann et al., 2015) in R v.3.6.0. Specifically, two ANOVAs were employed. The first ANOVA over midline electrodes included the factors Site (5 levels: Fz, FCz, Cz, CPz, Pz) and Condition. The factor Condition included three levels in Experiment 1 (acquisition phase; old pairs in the test phase; new pairs in the test phase) and four levels in Experiment 2 (acquisition phase; old pairs, new pairs, and old-new pairs in the test phase), see Figures 1 and 3 for traces of the ERP signals at midline electrodes.

**Figure 1.**
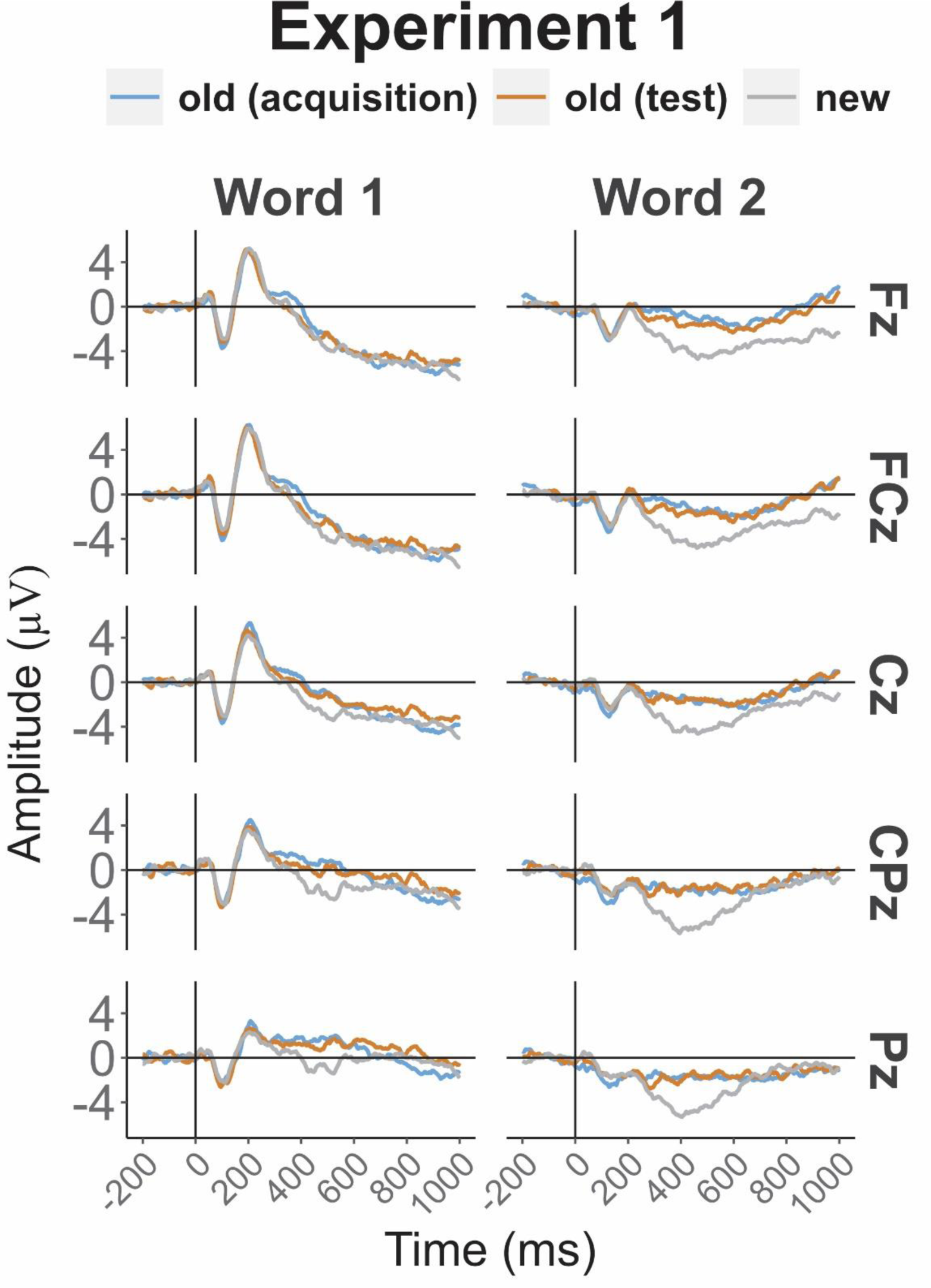
Midline channels ERP waveforms to the first and second words in pair in the first experiment.

**Figure 2.**
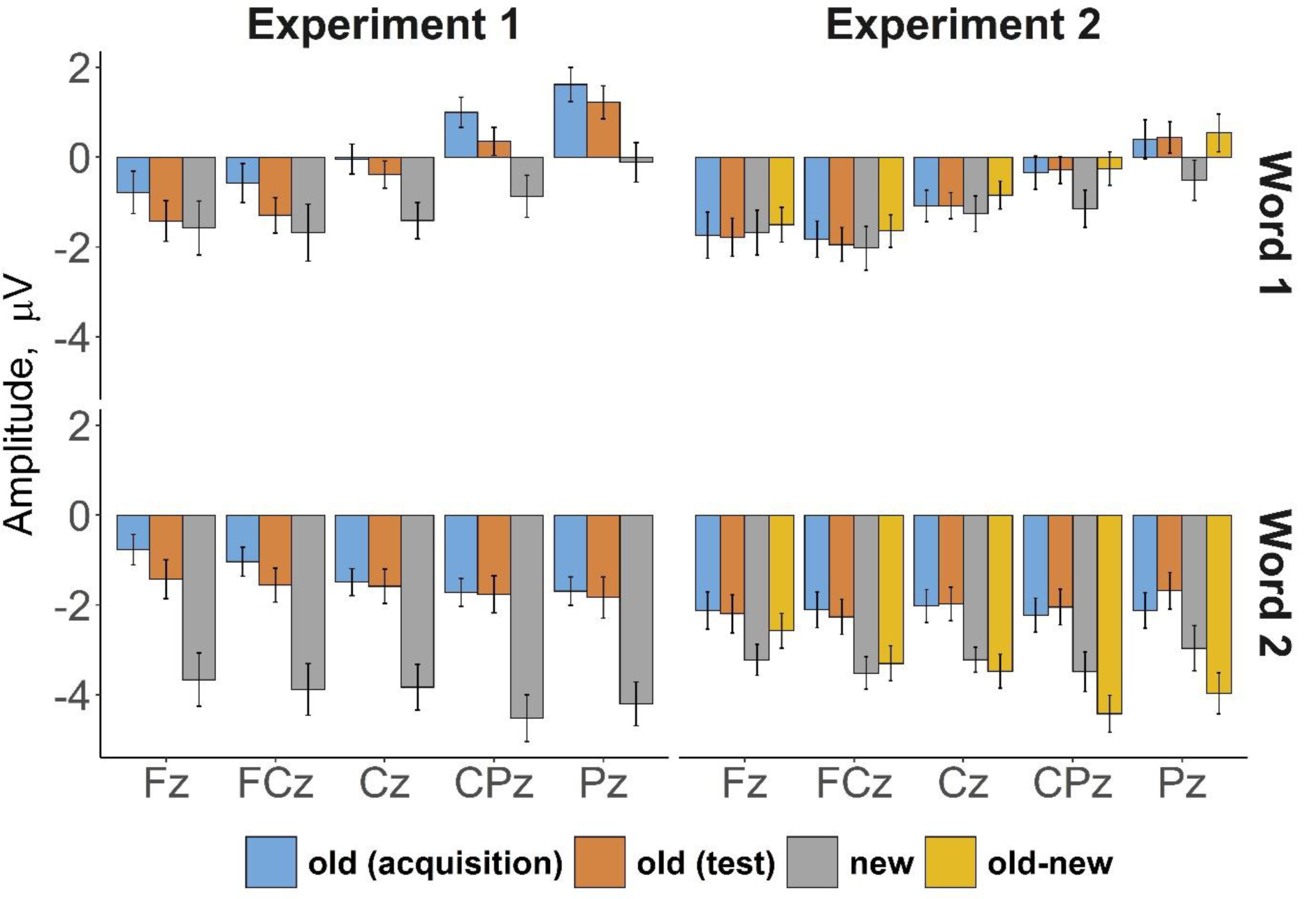
Averaged in N400 time window (250-500 ms) ERP amplitudes at midline channels. Error bars represent the standard error of the mean.

**Figure 3.**
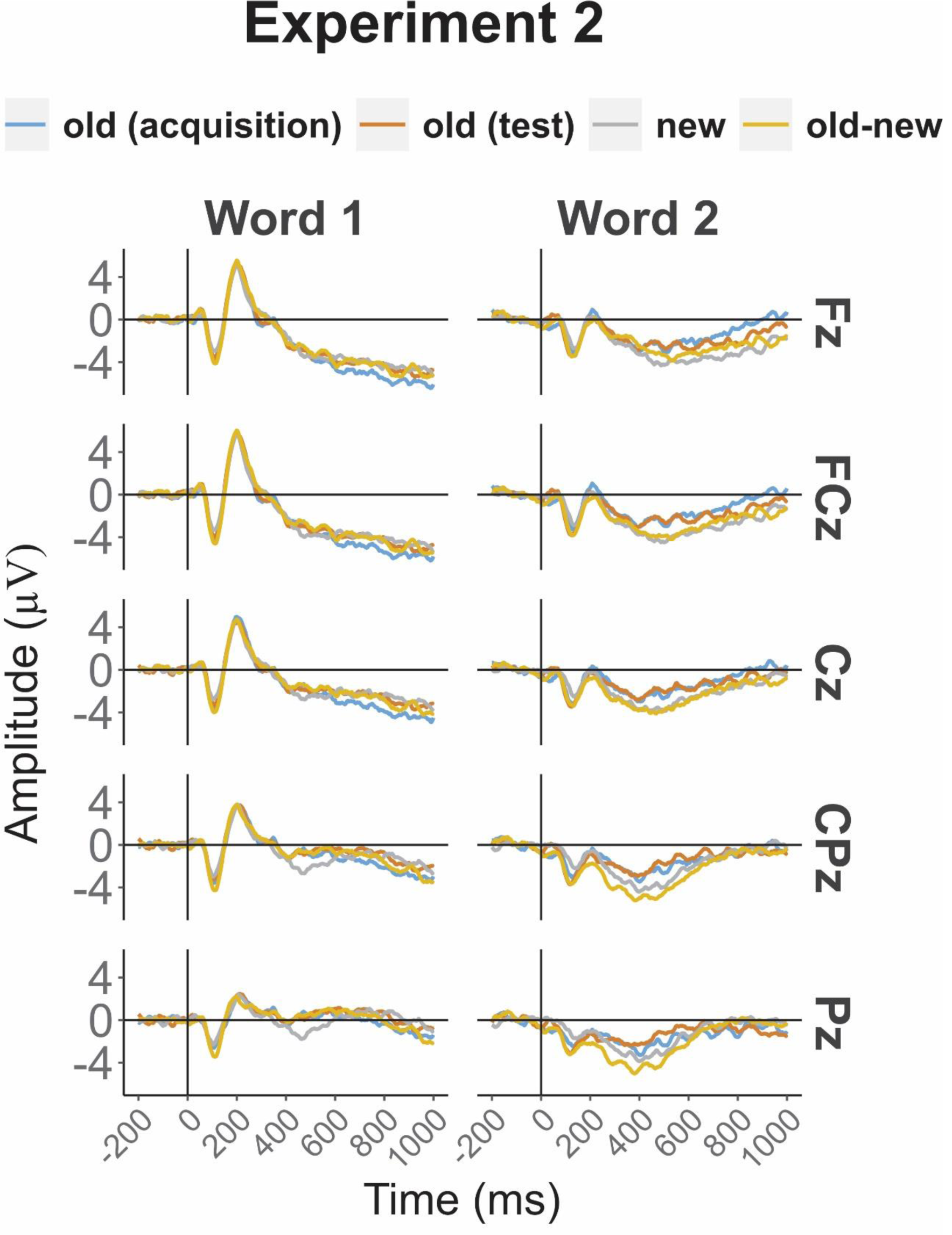
Midline channels ERP waveforms to the first and second words in pair in the second experiment.

The second ANOVA over lateral electrodes involved the factors Condition (identical to the first ANOVA), Region of Interest (ROI: 4 levels) and Side (2 levels, i.e. left versus right). For this sake we built average values for eight ROIs (4 ROIs on each side) including: left anterior ROI: F1, F3, FC1, FC3, FC5, FT7, left central ROI: C1, C3, C5, T7, left temporo-parietal ROI: CP1, CP3, CP5, TP7, P5, P7, left posterior ROI: P1, P3, PO3, PO7, O1, right anterior ROI: F2, F4, FC2, FC4, FC6, FT8, right central ROI: C2, C4, C6, T8, right temporo-parietal ROI: CP2, CP4, CP6, TP8, P6, P8 and right posterior ROI: P2, P4, PO4, PO8, O2.

The significance level was set at .05, and for all effects with more than two levels degrees of freedom were corrected for non-sphericity using Greenhouse-Geisser epsilon. Below, corrected degrees of freedom will be reported. Significant interactions of Condition with Site, ROI or Side were further investigated with post-hoc pair-wise t-tests using emmeans package in R (Lenth, Singmann, Love, 2018). Holm-corrected p-values (Holm, 1979) are reported in these post-hoc comparisons.

## 3. Results

### 3.1 Performance

In the first experiment, median reaction time (RT) was slower after new word pairs as compared to old pairs in the acquisition and test phases (*F*(2, 40) = 21.26, *p* < .001, *η*^2^ =.55), with means (averaged over individual *medians*) ± standard error (SE) being 937 ± 63.3 ms, 867 ± 62.7 ms, and 1120 ± 75.1 ms, for old (acquisition), old (test), and new pairs, resp. Pairwise comparisons confirmed the significance of the differences: old (acquisition) vs new, *p* < 0.001; old (test) vs new, *p* < 0.001; old (acquisition) vs old (test), *p* = 0.08.

The results of the second experiment were similar: old pairs in the acquisition and test phases were characterized by shorter RT than new and old-new pairs (F(2, 35) = 28.2, *p* < .001, *η*^2^=.57, means ± SE being 874 ± 59.7 ms (acquisition), 948 ± 49.3 ms (old pairs, test), 1090 ± 62 ms (new pairs), and 1140 ± 54.4 ms (old-new pairs). Pairwise comparisons revealed significant (all *p*s < .001) differences between old pairs (both acquisition and test), on the one hand, and new and old-new pairs, on the other hand. All other comparisons yielded *p*s > 0.4.

In Experiment 1 there was no difference between conditions in respect of the error rate (F(2, 32) = 0.74, p = .451, *η*^2^ = .03). In Experiment 2 the error rate was significantly higher after new second words than after old second words (F(2, 38) = 18.75, p < .001, *η*^2^ = .47). The absolute rate of errors was, however, very low: 3.4% in the old-new condition, 2.3% in the new-new condition, 0.4% in the old (test) condition, 0.8% in the acquisition, and, on average, 2.8% and 1.7% in the Experiment 1 and 2, respectively. Therefore, we did not exclude the few error trials from ERP averaging.

### 3.2 ERP

#### 3.2.1 Experiment I, First word

##### 3.2.1.1 N400 (250-550 ms time window)

###### Midline electrodes

As ERP waveforms in Figure 1 display (see also Figure 2), a significant main effect of Condition (*F*(1,30) = 6.24, *p* = .010, *η*^2^ =.23) resulted from the N400 amplitude for old pairs in the acquisition phase being significantly smaller than for new pairs (*p* = .028). Although the N400 was largest at Fz and FCz, between-condition differences at these sites did not reach significance (all *ps* > .05). At Cz, the N400 for new pairs was larger than in the acquisition phase (*p* = .025), and at Pz, it was larger for new pairs than for old pairs in both acquisition (*p* = .004) and test phase (*p* = .002). Finally, at CPz all three conditions differed significantly from each other: the N400 for old pairs was larger in the test phase than in the acquisition phase (*p* = .047), and for new pairs it was larger than for old pairs in both acquisition (*p* = .007) and test phase (*p* = .02). This pattern yielded a significant Condition by Site interaction: *F*(3,70) = 3.44, *p* = .018, *η*^2^ =.14.

###### Lateral electrodes

Like at the midline, the N400 for old pairs was significantly (*p* = .019) larger in the test phase than in the acquisition phase, leading to the main effect of Condition: *F*(1,30) = 7.06, *p* = .006, *η*^2^ =.25 (see Figure 4 for the spatial distribution of the effect).

**Figure 4.**
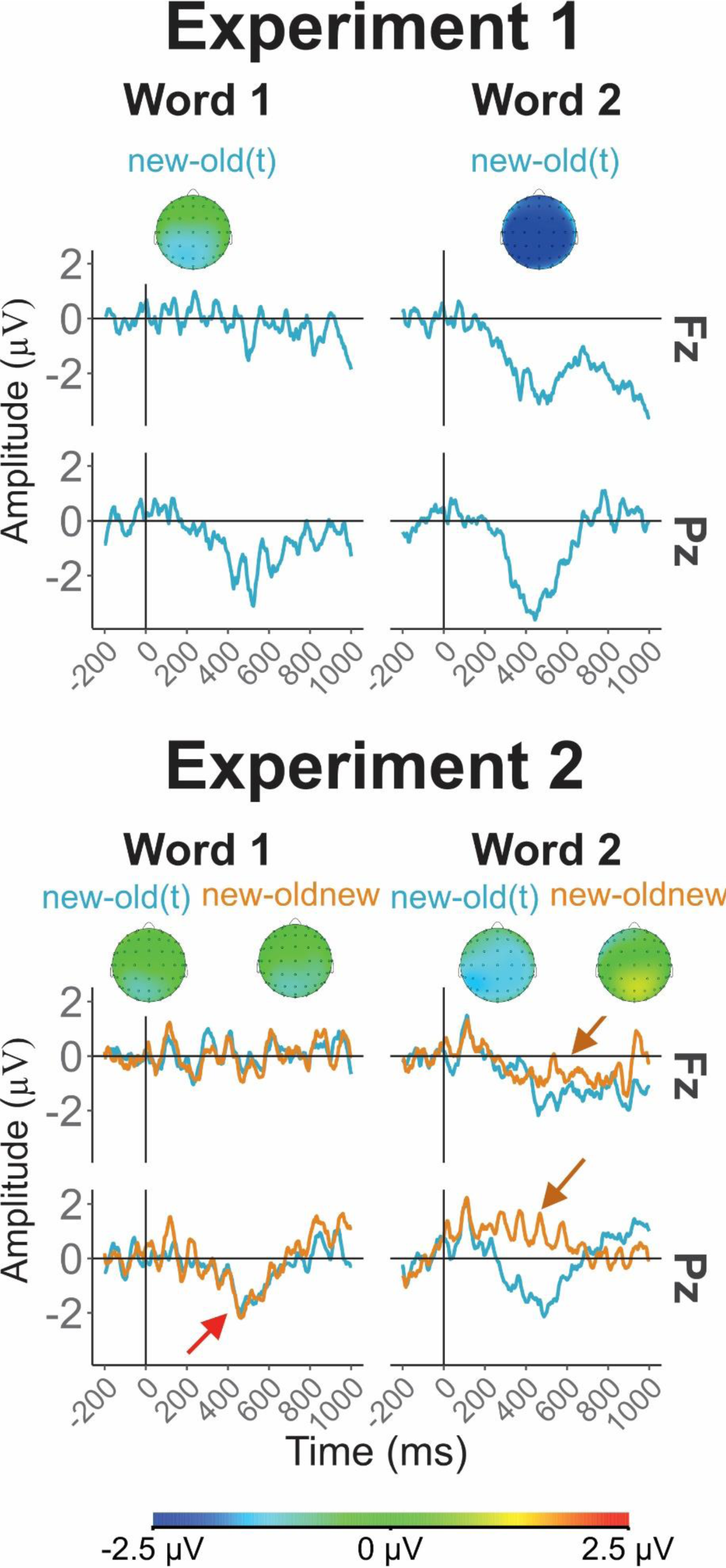
Difference ERP waveforms in Fz and Pz electrodes and difference topographic maps for N400 effect to the first and second words in both experiments. Notes: new-old(t) – the difference between new and old (test phase) word pairs; oldnew-new – the difference between old-new and new word pairs. Note that the difference between old and new FIRST words in Experiment 2 (**red arrow**) is a pure word repetition effect, because no effect of association within pairs can be expected. In contrast, the difference between SECOND words in old-new and new-new pairs in Experiment 2 (**orange arrows**) is a pure association effect, because the effect of repetition is identical in both compared conditions.

##### 3.2.1.2 LPC

No main effects of Condition or interactions including this factor were significant.

#### 3.2.2 Experiment I, Second word

##### 3.2.2.1 N400

###### Midline electrodes

As shown in Figure 1, the amplitude for new pairs was larger than the amplitude for old pairs in both test and acquisition phases (both *ps* < .001), yielding a significant main effect of Condition; *F*(1,39) = 15.12, *p* < .001, *η*^2^ = .42. The amplitude for old pairs did not differ between the acquisition phase and the test phase (corrected *p* =.538).

Generally N400 amplitude decreased in the posterior direction (main effect of Site: *F*(1,37) = 5.32, *p* = .012, *η*^2^ = .20). The effect of Condition, however, was not modulated by Site (*F*(3,71) = 1.94, *p* = .124, *η*^2^ = .08).

###### Lateral electrodes

A significant main effect of Condition (*F*(2,39) = 19.07, *p* < .001, *η*^2^ = .47) indicated that the amplitude for new pairs in the test phase was significantly larger than the amplitude for old pairs and in the acquisition phase (both *p* < .001). Like on the midline, the amplitude for old pairs did not differ between the acquisition and test phases (*p* = .53). None of the interactions of ROI and Side with Condition attained significance.

##### 3.2.2.2 LPC

###### Midline electrodes

The amplitude for new pairs in the test phase was smaller than the amplitude for old pairs in the test phase and in the acquisition phase creating a significant main effect of Condition (*F*(2,41) = 4.19, *p* =.023, *η*^*2*^ = .17). More specifically, the LPC was smaller for new pairs than for old pairs in the acquisition phase at Fz, FCz, and Cz (*p* = .002, .011, and .047, for Fz, FCz, and Cz, respectively). It was also smaller for new pairs than for old pairs in the test phase at the same sites (*p* = .002, .010, and .057, for Fz, FCz, and Cz, respectively). Accordingly, the main effect of Condition was significant at each of these locations (*F*(3,55) = 10.8, *p* <.001, *η*^*2*^ = .34; *F*(3,55) = 7.3, *p* =.002, *η*^2^ = .26; and *F*(3,55) = 4.6, *p* =.017, *η*^*2*^ = .18, for Fz, FCz and Cz, respectively). However, the difference between new and old pairs was not significant at posterior electrodes, which led to a highly significant Condition by Site interaction: *F*(3, 55) = 12.65, *p* <.001, *η*^*2*^ = .38.

###### Lateral electrodes

Like at the midline, the amplitude for new pairs in the test phase was smaller than old pairs in the acquisition phase giving rise to a significant main effect of Condition (*F*(2, 39) = 3.31, *p* = .05, *η*^*2*^ = .14). The effect of Condition was significant over anterior (*F*(2, 41) = 8.6, *p* <.001, *η*^*2*^ = .29) and central (*F*(2,36) = 5.8, *p* = .009, *η*^*2*^ = .22) but not in more posterior areas (Condition x ROI: *F*(2, 41) = 11.09, *p* <.001, *η*^*2*^ = .35). Pairwise comparisons within the ROIs revealed a pattern similar to the midline: the LPC was larger for old than for new pairs (old (acquisition) vs new: p = .005, .026 in anterior and central ROIs, respectively; old (test) vs new p = .007, .033 in anterior and central ROIs, respectively). Thus, the second word elicited a larger LPC in old pairs than in new pairs primarily at the anterior leads.

#### 3.2.3 Experiment II: First word

##### 3.2.3.1 N400

###### Midline electrodes

ERP waveforms at midline electrodes (Fz, FCz, Cz, CPz, and Pz) to the first word in the second experiment are shown in Figure 3. While the main effects of Condition was not significant (F(2,49) = 1.2, p = .3, *η*^*2*^ = .06), its interaction with Site was (F(4,83) = 4.3, p = .003, *η*^*2*^ = .18). Pairwise comparisons at each channel revealed that the effect was driven by a larger N400 to new first words as compared with old (test) first words at Pz (new vs. oldnew pairs: p = .018, new vs old pairs: p = .044).

###### Lateral electrodes

The main effect of Condition and its interactions with other factors were not significant. Generally, the amplitude of N400 was larger on the right than on the left side (main effect of Side: F(1,20) = 6.94, p = .016, *η*^*2*^ = .26). It was maximal over anterior ROIs and decreased in the posterior direction (main effect of ROI: F(1, 23) = 6.47, p =.015, *η*^*2*^ = .24).

##### 3.2.3.2 LPC

No interactions or main effects of Condition were found neither in the midline nor in the lateral electrodes.

#### 3.2.4 Experiment II, Second word

##### 3.2.4.1 N400

###### Midline electrodes

As can be seen in Figure 3 (and further illustrated in Figure 2), the amplitude to new second words was significantly larger than for old words in the test phase and in the acquisition phase (main effect of Condition: *F*(3,53) = 7.37, *p* < .001, *η*^2^ = .27; all pairwise *ps* < .05). In addition to the difference between new and old words, which was significant at Cz, CPz, and Pz, new words in old-new pairs elicited a significantly larger N400 than new words in new-new pairs at Pz (*p* = .039) and CPz (*p* = .018), thus resulting in a highly significant Condition by Site interaction: *F*(3, 60) = 6.38, *p* < .001, *η*^2^ = .24).

###### Lateral electrodes

Like at the midline, a significant main effect of Condition (*F*(3, 53) = 7.54, *p* < .001, *η*^2^ = .27) emerged from the fact that the amplitude for old-new pairs was significantly larger than for old pairs in the test phase (*p* < .001) and the acquisition phase (*p* = .007). The amplitude for new pairs was also significantly larger than for old pairs in the test phase (*p* = .001) and the acquisition phase (*p* = .007). The difference between old pairs in the acquisition phase and the test phase was not significant.

The N400 was larger over the left hemisphere (*F*(1,20) = 14.4, *p* = .001, *η*^2^ = .42). The difference between old and new second words was significant (all *p*s <. 05) in all ROIs except anterior ones (see Figures 2 and 3). The effect was stronger on the left side than on the right side. These findings manifested themselves in the significant interactions between Condition and ROI (*F*(3,52) = 5.24, *p* = .005, *η*^2^ = .21), between ROI and Side (*F*(2,36) = 5.82, *p* = .008, *η*^2^ = .23), as well as in a triple interaction of Condition, ROI, and Side (*F*(4,89) = 3.31, *p* = .01, *η*^2^ = .11). The differences between the acquisition phase and the same (i.e., old) stimuli in the test phase were not significant in any region.

##### 3.2.4.2 LPC

###### Midline electrodes

The amplitude was significantly larger in both old conditions (i.e., in acquisition and test) than in both new conditions (i.e., old-new and new-new) at Fz and FCz (local effects of Condition: *F*(2,45) = 4.96, *p* = .009, *η*^2^ = .19, and *F*(2,49) = 3.7, *p* = .025, *η*^2^ = .15, for Fz and FCz, respectively; all pairwise differences between old and new significant with *p*s < .05). There was no between-condition differences at more posterior sites, resulting in a highly significant Condition by Site interaction: *F*(3,51) = 7.37, *p* < .001, *η*^2^ = .27. The main effect of Condition was not significant.

###### Lateral electrodes

At lateral electrodes there was no significant main effect of Condition, but the interactions Condition by ROI (*F*(3,53) = 14.68, *p* < .001, *η*^2^ = .42) was significant. A further analysis showed that the main effect of Condition attained significance only in the anterior ROIs (*F*(2,50) = 5.6, *p* = .004, *η*^2^ = .22), where the LPC was larger in both old conditions as compared to new condition (both *p*s < .05) without significant differences in other comparisons.

## 4. Discussion

In the following, we shall first summarize the obtained results. Then, we shall discuss whether, and in what extent, the standard repetition effect could explain these results. After this, we shall discuss other factors necessary for explanation. Finally, we shall present some speculations about possible associative mechanisms and discuss limitations of our study.

### 4.1 Summary of the data

The study aimed to investigate the effect of unintentional learning of semantically unrelated word pairs on semantic ERP components. Two experiments were carried out, both containing an acquisition phase and a test phase. In both experiments, healthy subjects heard a series of semantically unrelated word pairs. A portion of the words were repetitions of previously presented items. In the test phase of the first experiment, “old” word pairs repeated word pairs presented during the acquisition phase, while “new” pairs contained other words that had not been presented before. In the test phase of the second experiment, a third condition was added in which the first word in a pair was one of the words presented during acquisition, but the second word was new.

The strongest effects were obtained in the time window of 250-550 ms, which we identified as the N400. In the *first experiment*, the amplitude was larger for new words than for old words. Although the effect was obtained in response to both words in the pair, it was stronger to second than to first words.

In the *second experiment*, the amplitude of the N400 was larger to new than old first words at posterior leads only (no main Condition effect, but Condition by Site interaction). The N400 was significantly larger to new second words than to old second words at all sites. The effect was most pronounced in the central, temporo-parietal and posterior regions. At these locations, furthermore, new second words elicited a larger N400 when the first word in the pair was old, as compared with the condition when the first word was also new.

In *both experiments*, additionally, the LPC (time window 550 - 1000 ms^1^) at frontal leads had larger amplitude to old second words than to new second words. The effect was absent to first words.

### 4.2 ERP repetition effect

Rugg (1985) was probably the first who examined the effects of repetition on the ERPs elicited by words in lists. He compared the effects of repetition to those of associative semantic priming and found that both repeated and semantically primed words elicited more positive ERPs than did new or unrelated words in the latency range of 300 to 500 msec post-stimulus. This was consistent with the conclusion that both repeated and related words elicit a smaller N400 in lists.

In the following study, Rugg (1987) investigated a combination of repetition and semantic effects. Unlike our study, the experiments included both words and pronounceable non-words. Subjects silently counted occasional non-words against a background of meaningful words, a portion of which were either semantic associates or repetitions of a preceding word. Repeated words were distinguished by waveforms containing an initial transient negative-going deflection called N140. In contrast to the current study, the repetition effect in Rugg (1987) did not appear as a typical N400. Since that experiment included two types of stimuli (words and non-words), onset latency differed between the two studies. One may suppose (although we do not have a direct comparison) that the task to distinguish words from non-words is more difficult, or stronger interfere with semantic processing, than our task to count syllables, and that for this reason word/non-word discrimination took more time. In the light of the above, it seems reasonable to assume that the differences between word and non-word repetition effects on ERPs are attributable to the dissimilar properties of these two types of item, which is completely different from our experiment restricted to one kind of stimulus, unrelated word pairs.

Van Petten, Kutas, Kluender, Mitchiner and McIsaac (1991) additionally investigated the effects of repetition as a function of word frequency. Whereas repeated low-frequency words elicited a positive ERP component after 500 ms post-stimulus, the predominant effect for repeated high-frequency words was a reduction of N400 amplitude. The current result is in line with that finding because most words presented in our study belonged to highly to moderately frequent German words.

Several studies showed that the difference between ERPs to new and repeated words begins only after N400 and mainly covers the time window roughly corresponding to the LPC in the present study (e.g., Curran, 1999; Balass et al., 2010). In other studies, the effect starts about 300 ms post stimulus but attains its maximum at about 600 ms (e.g., Rugg et al., 1994, 1997; van Strien et al., 2007). Most frequently, however, the repetition effect involves both N400 and LPC time windows (e.g., Karayanidis et al., 1991; Kutas & Van Petten, 1988; Fischler & Raney, 1991; Race et al., 2010). The earlier effect, that can be regarded as N400 attenuation to repeated words, usually has a parietal maximum. The spatial distribution of the later effect (larger LPC positivity to repeated words) is more variable and has, to our best knowledge, not been an object of a special analysis. Most studies clearly show that this effect was also strongest in parietal areas (e.g., Rodriguez-Fornells et al 2002; Swick, 1998; Van Strien et al., 2007; Race et al., 2010; Olichney et al., 2006); therefore, it appeared to be a direct continuation of the N400 effect. However, other publications indicate a maximal LPC effect at frontal leads (e.g., Bentin & Peled, 1990, nonword detection task; Besson et al., 1992), whereas Rugg et al. (1997) observed its maximum at Cz decreasing in both frontal and parietal direction.

### 4.3 Repetition effect versus learnt association

Contrary to the previous studies, we presented words not in lists (e.g., Rugg, 1985, 1987; Rugg et al., 1994, 1997) and not in sentences (e.g., Besson et al., 1992; Calloway & Perfetti, 2019) but in pairs. Nevertheless, our experiments involved massive repetition of the “old” words, and thus repetition effects were expected. As already said above, repetition effect is typically strong after 500 ms, corresponding to the LPC. The similar effect in our experiments was observed only in fronto-central areas and only to the second words in pairs. The former finding does not contradict the repetition effect interpretation; as stated above, most electrophysiological word repetition studies showed a posterior LPC effect, but a minority of studies found fronto-central effects as well. The latter finding appears more surprising. Both first and second “old” words were repeated with the same frequency, thus the repetition effect should be equally expected for both. Even if the rather short and variable onset-to-onset interval within pairs might substantially suppress the LPC effect to the first words, this can only be a partial explanation. This factor can hardly explain the fact that first words not only elicited no significant LPC effect (i.e., no difference between old and new words), but no LPC altogether.

Turning now to the dynamics of the N400, we see a scale of effects. The strongest increase of the N400 was elicited by new second words in Experiment 2 in the pairs in which the first word was old. This increase was significantly larger than the corresponding increase to the new second words in the pairs in which the first word was also new. Much weaker but still significant at most sites was N400 increment to new (as compared with old) first words in Experiment 1. Finally, the weakest change was N400 increment to new first words in Experiment 2, which was significant only at posterior electrodes.

To make an interim conclusion, both N400 and LPC dynamics are rather unusual for the typical repetition effect. Already their different topography indicates the presence of at least two distinct mechanisms. Additionally, a typical repetition effect might manifest itself in N400 decrease to old words in test compared with the same words during acquisition, but the tendency was rather opposite. In Experiment 1 the N400 was even significantly larger to old words in test than acquisition, but this significance might be spurious because it was not replicated in Experiment 2. Anyway, however, the N400 was definitely *not smaller* to old words in test than acquisition as the typical repetition effect would suggest.

If word repetition was the dominant effect in the N400 time window, (i) the response to the first word would not depend on the second word, and (ii) vice versa, the response to the second word would not depend on the first one. As regards the statement (ii), it is definitely wrong. First, the increase of the N400 to new words was about twice as strong for second words as for the first words. Additionally, at posterior sites in Experiment 2, the increase of N400 amplitude to the new second word was significantly larger when the preceding word was old than when it was also new.

As regards the statement (i), the evidence is weaker. Although the N400 effect to the first word was significant at almost all sites in Experiment 1 but only over parietal cortex in Experiment 2, and although the size of this effect was (except for parietal and occipital areas) almost twice larger in Experiment 1 than Experiment 2, the interactions Condition by Experiment and Condition by Site (or ROI) by Experiment were not significant. Remember that in Experiment 1 an old first word always indicated that the second word would also be old, i.e., the whole pair is old. In Experiment 2, in contrast, after an old first word participants could not know, whether the presented pair is old or new. Therefore, if the difference between the two Experiments is real, it is probably due to the repetition of the whole pair being an additional factor distinct from word repetition. In this case the statement (i) would also be wrong, but a definite decision can only be made after a replication experiment.

To summarize, the data indicate the presence of at least three different effects. Two of them (the parietal N400 decrement to new first words in Experiment 2 and the fronto-central LPC increment to new second word) may be attributed to the same factor of word repetition, but differ so much from each other that they cannot be regarded as a single “repetition effect”. The concept of several functionally distinct electrophysiological repetition effects has been suggested in the literature (Guillem et al., 1999; Penny et al., 2001; Race et al., 2010; Calloway & Perfetti, 2019), but none of the proposed models can be directly applied to the present data due to the large variety in experimental designs. Over and above the two repetition effects, the third effect reflects the specifics of the presented stimulus material, i.e., the relations between words in pairs.

### 4.4 Within-pair associations

Therefore, a considerable portion of the N400 effect distinguishing between the responses to learnt and new word pairs cannot be attributed to word repetition but is due to a relation emerging between the members of a repeated pair. There is a long lasting discussion in the N400 literature on whether, or to what extent, this component manifests automatic versus controlled processes involved in building semantic associations. The former processes are theorized on the basis of the Automatic Spreading Activation (*ASA)* model (Collins & Loftus, 1975). According to many authors (e.g., Neely, 1977; Neely & Keefe, 1989; Hill, Strube, Roesch-Ely & Weisbrod, 2002), the mental lexicon is assumed to be a semantic network with related words in neighboring nodes. ASA occurs because the lexical access to a word activates the corresponding node and this activation spreads to the neighboring nodes corresponding to related words. When a prime is presented, a set of nodes corresponding to related words are pre-activated. The spreading activation lowers the threshold for the processing of all related nodes. Therefore, when a related (i.e., already pre-activated) word is presented, its processing requires *less* additional activation, thus resulting in a *smaller* amplitude of the N400 to this word. In accordance with this idea, the semantic facilitation effect in a lexical decision task is usually attributed to the spreading-activation process (e.g., Schvaneveldt & Meyer, 1973; Koriat, 1981). Note that the ASA model presumes the same basic underlying mechanism for N400 attenuation to *semantically related* second words and to second words in word pairs that have been recently presented and learnt *in semantically unrelated sets*, as compared with semantically unrelated word pairs *presented for the first time*.

Esper (1973; cit. for Maki, McKinley, & Thompson, 2004) emphasized the principal difference between two kinds of associations: inner associations are based on semantic or logical relationships and not on empirical frequencies. Thus CAT and MAMMAL are internally associated even though these two words rarely occur close in most real sentences. In contrast, CAT and DOG are, to a large extent, an outer association because the two words frequently co-occur.

The process of building such associations between previously unrelated words can be conceived of as a chain of mediated priming processes whose ERP correlates were studied by Chwilla et al. (2000) and Chwilla & Kolk (2002). The former study investigated mediated two-step priming with one intervening concept (e.g., the prime LION activates the target STRIPES via TIGER). The latter study showed that the N400 to the target can be suppressed even when the prime is separated by several mediators (e.g., a prime MANE is associated with the target STRIPES through mediators LION and TIGER).

Following these findings, we assume that, when an unrelated word pair (e.g., HORSE – FAST) is repeated, the brain does not just notice the empirical co-occurrence of the two stimuli. Rather, it is searching for an interpretation of this newly emerging connection (see Kotchoubey, 2006; Schlesewsky & Bornkessel-Schlesewski, 2019). The activation spreads to broader and broader networks of nodes related to the two words. Some of these activated nodes are related to *both* words in the pair and, under their activation, build a semantic bridge between them. In the above example, repeated presentation of HORSE may activate such neighboring nodes as LEGS, RIDING, and finally RACE, which should run FAST to win a match. Thus an association that is primarily built as a purely outer one (in the sense of Esper, 1973), can acquire inner properties.

The automaticity of the processes underlying the N400 effect can, however, be questioned. Several studies demonstrated an automatic N400 effect by using sensory masking, i.e., non-masked target words were presented after strongly masked (unrecognizable) primes (e.g., Dehaene et al., 2001; Kiefer, 2002; Misra & Holcomb, 2003). Daltrozzo et al. (2012) used the same masking approach but masked not only primes but also targets. At the strongest level of masking, participants were unable to distinguish between congruent and incongruent conditions (i.e., the error rate was about 50%), but still could correctly compare the completely masked target word with a subsequent probe word presented without a mask. No N400 to incongruent targets was found in this conditions, while the LPC disappeared at even lighter masking levels when the error rate was still very low (about 15%). These findings contradict the results of the experiments in which only primes were masked indicating that the N400 effect cannot depend only on ASA, because the effect can disappear when stimuli are not explicitly processed, even if their implicit (i.e., automatic) semantic processing can take place.

We believe that the same is true for the present data. A completely automatic activation would result in a decrease of the N400 to repeated (i.e., associated) words, thus explaining why the N400 effect was so much stronger to the second than to the first word. It cannot, however, explain an additional N400 increase to the second word in old-new pairs as compared with new-new pairs. Actually, the ASA model predicts only N400 decrement to primed words but not an increment to non-primed words, which is in line with the observation that an N400 is usually present to all non-primed words (e.g., Van Petten, 1995). In the old-new versus new-new comparison, however, both classes of second words are non-primed, and there is no reason for an N400 difference between them.

This difference can, from our point of view, be only explained by an active anticipatory process emerging within a learnt pair. This process can be designated as, or related to, semantic integration (e.g., van Berkum et al., 1999; Sitnikova et al., 2003), reallocation of cognitive resources necessary for such integration (e.g., Kotchoubey, 2006; Bornkessel-Schlesewski & Schlesewsky, 2019), or expectancy developing after the first word (if the term “expectancy” is used in a broad sense including unconscious and unintentional preparation). When the first word in a pair in Experiment 2 was new, the following word was also new, and there was no need to look for a meaningful relation between two novel, non-learnt, non-associated stimuli. When, on the other hand, the first word was old, a previously learnt association was activated. On the background of this activation, an unexpected second word led to redirection of resources to restore the “lost agreement” between the two members of the pair.

### 4.5. Limitations and further possibilities

Although numerous data of the literature suggest that the effects should be specific for meaningful words, or maybe for meaningful stimuli in general (Sitnikova et al., 2003), this supposed specificity is not warranted. Perhaps we need at least two additional experiments in which (i) meaningless non-words and (ii) equally complex (acoustically) but not word-like sounds would be presented in pairs like in the current experiments.

Another unclear issue concerns the role of attention. The ASA model presumes that the effects should be largely independent of attention. Some data indicate that active instruction to respond to stimuli can increase the amplitude of the N400 as compared with more passive conditions (e.g., Erlbeck et al., 2014). Particularly, the process of semantic integration supposedly responsible for the increase of the parietal N400 in old-new pairs, can be attention-dependent. In the present study, we used an instruction that demanded participants’ attention to stimuli. In some contexts, and specifically in clinical applications of the paradigm on patients with preserved semantic abilities but attentional problems, the issue of attention may be very important. A replication of the present experiments without active instruction would help to answer the corresponding questions.

The problem of attention is closely related to that of consciousness. Earlier studies indicating that some N400 effects can be obtained without conscious awareness of stimuli (e.g., Dehaene et al., 2001; Kiefer, 2002) were questioned when stronger control of the lack of awareness resulted in the disappearance of the N400 (Daltrozzo et al., 2012). Particularly, the hypothesis that presentation of the first word of a learnt word pair activates “expectancy” of the second word, leads to the question as to which extent the expectancy is a conscious process. Further experiments varying both perceivability of stimuli and the task (e.g., the relevant task to discriminate between new and old pairs instead of the irrelevant task used in the current study) are necessary to explore this issue.

## 5. Conclusions

We found that the acquisition of new word association in semantically unrelated word pairs results in a change of N400 amplitude very much similar to that observed in semantically related word pairs. The mere effect of word repetition, though present, does not explain all obtained results. Consistent co-occurrence of words elicited additional effects more specifically related to associative learning. The relative role of automatic processes (particularly, the Automatic Spreading Activation) and controlled, potentially attention-dependent processes (e.g., semantic integration) should be investigated in further experiments.

## Acknowledgments

The study was supported by the German Research Society (Deutsche Forschungsgemeinschaft), Grant KO‐1753/13.

## Footnotes

1 Because of the lack of a clear LPC peak, we also tried other window definitions of this component. The result was always the same.

## Notes

### Competing Interest Statement

The authors have declared no competing interest.

